# Enlarging the Scope of Randomization and Permutation Tests in Neuroimaging and Neuroscience

**DOI:** 10.1101/685560

**Authors:** Eric Maris

## Abstract

Especially for the high-dimensional data collected in neuroscience, nonparametric statistical tests are an excellent alternative for parametric statistical tests. Because of the freedom to use any function of the data as a test statistic, nonparametric tests have the potential for a drastic increase in sensitivity by making a biologically-informed choice for a test statistic. In a companion paper (Geerligs & Maris, 2020), we demonstrate that such a drastic increase is actually possible. This increase in sensitivity is only useful if, at the same time, the false alarm (FA) rate can be controlled. However, for some study types (e.g., within-participant studies), nonparametric tests do not control the FA rate (see Eklund, Nichols, & Knutsson, 2016). In the present paper, we present a family of nonparametric randomization and permutation tests of which we prove exact FA rate control. Crucially, these proofs hold for a much larger family of study types than before, and they include both within-participant studies and studies in which the explanatory variable is not under experimental control. The crucial element of this statistical innovation is the adoption of a novel but highly relevant null hypothesis: statistical independence between the biological and the explanatory variable.

## Introduction

Nonparametric statistical tests are an excellent alternative for several parametric statistical tests, especially when analyzing the high-dimensional biological data that are typically collected in cognitive and medical neuroscience. This is for two reasons: (1) their false alarm rate control does not depend on auxiliary assumptions or asymptotic arguments, and (2) every function of the data can be used as a test statistic, which creates the opportunity for a drastic increase in sensitivity. Especially the first reason has received a lot of attention, and this is because the probability distribution of the biological data often violates the auxiliary assumptions of the normal theory framework that underlies parametric statistical tests. Violation of these auxiliary assumptions often results in an inflated false alarm (FA) rate (the probability of falsely rejecting the null hypothesis). This has recently been demonstrated in a large-scale simulation study using data of published neuroimaging studies (Eklund et al., 2016).

The main motivation for the present paper is the potential for a drastic increase in sensitivity by using a test statistic that incorporates prior knowledge about the signal (the effect) and the noise between which one wants to discriminate. In a companion paper (Geerligs & Maris, 2020), we demonstrate that such a drastic increase is actually possible. Specifically, Geerligs and Maris (2020) introduce the so-called min(p) method for combining a set of basic test statistics with different sensitivity profiles (e.g., narrow strong effects and widespread weak effects). This min(p) method produces a single test statistic with a much broader sensitivity profile, encompassing the sensitivity profiles of the more specialized basic test statistics. In a simulation study, Geerligs and Maris (2020) demonstrated that a test statistic can be produced with a much higher sensitivity than the sensitivities of the existing methods for the statistical analysis of fMRI data.

Crucially, an increase in sensitivity is only useful if, at the same time, the specificity is controlled. Specificity is typically quantified as the FA rate. Unfortunately, FA rate control is not always guaranteed for a nonparametric test. For example, in the simulation study by Eklund et al. (2016), a permutation test for a within-participants design (called “one-sample” test by Eklund et al., 2016) did not control the FA rate. However, in the same simulation study, it was also shown that a permutation test for a between-participants design (called “two-sample” test by Eklund et al., 2016) did control the FA rate.

Besides the failure of the permutation test to control the FA rate for all study designs, Eklund et al. (2016) also demonstrated that different preprocessing pipelines (corresponding to different default options in the major fMRI analysis packages) result in very different empirical FA rates. Thus, the empirical FA rate not only depends on standard settings, such as the cluster-forming threshold (which can be set by all packages), but also on settings that cannot be controlled in at least some of the packages (Bowring, Maumet, & Nichols, 2018). This state of affairs is worrisome, especially in times when scientists experience the pressure to demonstrate the replicability of their results.

In the present paper, we demonstrate that nonparametric statistical tests have a much larger scope than is often assumed. Specifically, we describe a nonparametric test that controls the FA rate for a wide range of study designs, including the within-participants design for which the one-sample test of Eklund et al. (2016) did not achieve this. In addition, the FA rate is also controlled for studies in which the explanatory variable (see further) is not under experimental control. To demonstrate this FA rate control, it is necessary to introduce a novel but highly relevant null hypothesis: *statistical independence between the biological and the explanatory variable*. The terms “biological” and “explanatory variable” require some explanation. In most papers and handbooks, instead of “biological” and “explanatory variable”, the authors use the terms “dependent” and “independent variable”. However, these terms would create too much semantic overlap with the term “statistical independence”. The terms “biological” and “explanatory variable” have the advantage that their meaning is very close to our target field of application: neuroimaging and neuroscience.

In the remainder of this paper, we first give a brief overview of the differences between parametric and nonparametric statistical tests. Next, we describe how to perform a randomization test for random effects in studies with a within-participants manipulation of the explanatory variable. This description serves as a motivation for the remainder of the paper, which is more formal. This more formal part is divided in three sections:

1. Randomization tests for studies with a between-units manipulation of the explanatory variable
2. Randomization tests for studies with a within-participants manipulation of the explanatory variable
3. Permutation tests for studies in which the explanatory variable is not under experimental control (which we call a “natural explanatory variable” in the following).

## Parametric versus Nonparametric Statistical Tests

Below, is a four-point schema that specifies how a statistical test is performed. This schema is valid for both parametric and nonparametric statistical tests:

1. Evaluate some test statistic that is a function of both the biological and the explanatory variable. For example, in an independent samples t-statistic, the participant-specific average magnetic resonance (MR) signal in a given voxel could be the biological variable, and the participant’s membership to some group (e.g., patient or control, old or young) could be the explanatory variable.
2. Derive or construct a probability distribution of this test statistic under some null hypothesis. This distribution will be called the *reference distribution*.
3. Calculate the probability under the reference distribution of observing a test statistic that is more extreme than the observed value of the test statistic. This probability is typically called the *p-value*.
4. Reject the hypothesis of statistical independence if that p-value is less than some nominal value (typically, 0.05 or 0.01).

*Valid* statistical tests must control the false alarm (FA) rate. That is, the probability of falsely accepting the null hypothesis must be less than the nominal value mentioned in point 4. *Good* statistical tests must be valid and also have a high power (sensitivity), that is, a high probability of rejecting the null hypothesis when it is in fact false.

Nonparametric statistical tests differ from the parametric ones in two aspects. First, valid nonparametric tests can be constructed for arbitrary test statistics, and therefore are not limited to the family of tests statistics that has been derived under the normal distribution (T-test, F-test, …). This freedom creates the possibility to incorporate biophysically motivated constraints in the test statistic, such as clustering over the spatial, temporal, and spectral dimensions of the data (Maris & Oostenveld, 2007). Additionally, this freedom also creates the opportunity to combine test statistics with different sensitivity profiles (e.g., cluster-based tests with different cluster-defining thresholds), which increases the width of the sensitivity spectrum. This is demonstrated in a companion paper (Geerligs and Maris, 2019).

A second important difference with the parametric statistical tests pertains to the reference distribution. In the parametric framework, the reference distribution of a test statistic is *derived* from assumptions about the data whereas in the nonparametric framework it is *constructed* from the observed data. In the present paper, we distinguish between randomization and permutation tests. The reference distribution for a randomization test is called the *randomization distribution*, and the one for a permutation test is called the *permutation distribution*. A randomization test requires that the units (participants, event times, …) are assigned to the experimental conditions on the basis of a randomization mechanism (typically, calls to a pseudo-random number generator). Thus, a randomization test involves an explanatory variable that is under experimental control. This is not necessarily the case for a permutation test, which can also be applied to a natural explanatory variable (e.g., gender, disease status, age, accuracy), of which the values are observed instead of assigned.

Randomization and permutation tests can be motivated in different ways (Maris & Oostenveld, 2007; Pesarin, 2001; Raz, Zheng, Ombao, & Turetsky, 2003; Winkler, Ridgway, Webster, Smith, & Nichols, 2014). In this paper, we will motivate them from the perspective of testing the null hypothesis of statistical independence between the biological and the explanatory variable. Specifically, we will prove that a decision on the basis of a randomization or permutation p-value controls the FA rate under this null hypothesis. We will present our main theoretical results (pertaining to FA rate control) first for randomization tests. Only in the last section, we will describe how these results can be generalized to a particular class of permutation tests, which are specific for natural explanatory variables. This class of permutation tests partially overlaps with the popular existing permutation tests for neuroimaging data (Eklund et al., 2016; Nichols & Holmes, 2002), but they are not identical. These popular existing permutation tests will be denoted as “sign-flipping tests”. Sign-flipping tests are motivated by the null hypothesis of symmetric error distributions (Nichols & Holmes, 2002; Winkler et al., 2014). As will be outlined in the following, this differs from the motivation of the randomization and permutation tests that we will introduce.

## A Randomization Test Alternative for the Sign-Flipping Test

This section serves as a motivating example. Specifically, we will demonstrate how to perform a statistical test that controls the FA rate for a study type for which this has not been demonstrated before. The formal proof of this FA rate control will be given later.

### The Design of the Study

We consider a multi-participant study in which every participant is observed in all experimental conditions of interest. These experimental conditions are the levels of the explanatory variable, and therefore we denote this study type as one in which the explanatory variable is manipulated *within* participants. As always, the research question pertains to differences between the experimental conditions in the measured biological variable. There are several variants of this study type, and the most typical ones are the trial-based (also called “blocked design”) and the event-based (also called “event-related design”) variant. For the present paper, the difference between these variants is not crucial, and therefore we will use “event time” both to denote the start of a trial (which lasts for several scans) and a single scan.

For the present paper, it is important to be explicit about how the event times are assigned to the experimental conditions (A and B). In most studies, the researcher makes a deliberate choice about the number of event times that must be assigned to each of the experimental conditions, and this number usually is chosen to be equal for each of these conditions. Next, the order of the experimental conditions across the event times is determined. There may be a single order that is used for all participants (e.g., AABBABAB), or there may be multiple orders (e.g., AABBABAB and BBAABABA). In the latter case, there are two ways to assign the participants to one of these orders: (1) at random (typically, by calling a pseudorandom number generator), or (2) on the basis of the researcher’s judgment (e.g., alternating the one and the other condition order). For the randomization test that will be introduced in the following, it is necessary that there are multiple orders and that the participants are assigned at random to one of them.

It is possible that one of the two conditions (say, B) is a so-called null event. A null event corresponds to the absence of some event of interest A (typically, a stimulus). A statistical test that is sensitive to the difference between A and the null event B, is called “testing against zero”. For the randomization test that will be introduced in the following, testing against zero is performed by comparing two condition orders (e.g., AABBABAB and BBAABABA) in which one condition corresponds to a null event.

The scenario described above involves a single set of event times for all participants and two possible condition orders to which a participant can be assigned. This is only the simplest of the scenarios for which our method is valid. In general, what is required for the validity of our method is that, for every participant, the event times can be partitioned into two or more sets, each of which corresponds to one condition. These event times and their partitioning may be different for different participants, resulting in different participant-specific sets of possible condition orders. We will return to this in a later section (*Randomization Tests for Studies with a Within-Participants Manipulation of the Explanatory Variable*).

### The Existing Statistical Tests

The existing statistical tests for this study type involve two steps: in the first step, the effect is quantified separately for every participant, and in the second step, these participant-specific effect quantifications are combined in a test statistic for which a p-value is calculated. There are many ways to quantify an effect. Nowadays, this quantification typically involves a participant-specific general linear model (GLM) analysis. This analysis returns condition-specific regression coefficients for the regressors that model the hemodynamic responses evoked by the events of interest. The effect quantification then typically is taken as the difference (contrast) between these condition-specific regression coefficients. Note that one of the two experimental conditions can also correspond to a null event. In the existing statistical tests, this null event may or may not be included in the effect quantification (by including a null event regressor in the GLM). As will be described later, in the statistical tests we propose, the null event is always included in the effect quantification.

With more than two experimental conditions, the effect quantification typically takes the form of an F-statistic, which depends on the ratio between the between-condition and the within-condition variance. The difference between two and more experimental conditions is irrelevant for our comparison of the parametric and the different nonparametric statistical tests on which we focus in this section. Therefore, we will only consider studies with two experimental conditions.

Next, a statistical test is performed, using as ingredients the participant-specific effect quantifications. In the parametric framework, this statistical test usually is a one-sample t-test, which tests the null hypothesis that the expected value of the effect quantification equals zero. This one-sample test is also called a paired- or a dependent samples t-test, in which “paired” and “dependent” refers to the fact that every participant is observed in two conditions. In the nonparametric framework, one usually performs a sign-flipping test (Nichols & Holmes, 2002; Winkler et al., 2014). The reference distribution of this test is obtained by randomly flipping the signs of the participant-specific effect quantifications. Both the parametric and the nonparametric statistical tests have their problems, and these will now be discussed.

### The Problems with Parametric Statistical Inference

It is well known that the FA rate control of a parametric statistical test depends on so-called auxiliary assumptions. For the one-sample t-test, these auxiliary assumptions are normality and between-participants statistical independence of the difference scores. (As an aside, all statistical tests, both parametric and non-parametric, assume between-participants statistical independence, and therefore we will not explicitly mention this auxiliary assumption anymore.) Fortunately, as a result of the central limit theorem, parametric statistical tests provide asymptotic control of the FA rate: with increasing sample size, their FA rate approaches the nominal alpha level. Although there are no objective guidelines for the required sample size to achieve a particular degree of FA rate control, it is often stated that one-sample t-test is robust against deviations from normality.

The situation is more serious when, instead of scalar observations, high-dimensional data are observed. An MR scan produces one signal per voxel, of which there are several thousands. Because there is one statistic per voxel-specific signal, there is a huge multiple comparisons problem (MCP). A common way to deal with this MCP is by calculating a function of these voxel-specific statistics, and using the function value as the actual test statistic on which the statistical decision (accept or reject the null hypothesis) is based. Popular functions are the maximum and the minimum, as well as functions that involve thresholding and clustering, such as the maximum cluster size and the maximum cluster sum.

In the parametric framework, these test statistics are then evaluated under reference distributions that are derived from the theory of Gaussian (or Student T) random fields (Friston et al., 1994). FA rate control using these reference distributions is only asymptotic. Specifically, for cluster-based statistical tests, only with an increasing cluster-forming threshold, the empirical FA rate approaches the nominal alpha level. By means of simulations, Eklund et al. (2016) demonstrated that the cluster-forming thresholds corresponding to voxel-level FA rates of 0.05 and 0.01 resulted in an unacceptably high empirical FA rate.

### The Problems with Nonparametric Statistical Inference

It is often claimed that nonparametric tests allow for testing a null hypothesis without relying on auxiliary assumptions. However, it is not always clear what is actually meant by this claim. Ideally, it means that, by means of a formal (mathematical) proof, it is shown that the FA rate of the statistical test is controlled under a null hypothesis that makes no assumptions about the parametric shape of the probability distribution that generates the data. Although there is theoretical work on the FA rate control of the sign-flipping test starting from the null hypothesis of symmetric error distributions (Nichols & Holmes, 2002; Winkler et al., 2014), no such formal proof has been given for this particular test. Therefore, statements about its FA rate control are typically based on simulation studies, such as those by Eklund et al. (2016). These authors demonstrated that the sign-flipping test in general failed to control the FA rate: depending on the dataset examined and the test statistic that was used (cluster- or non-cluster-based), the empirical (i.e., simulation-based) FA rate varied between 0.8 percent (too conservative) and 40 percent (too liberal). Eklund et al. (2016) attributed this failure to the asymmetry of the error distributions.

However, for some nonparametric tests, the empirical FA rate *is* controlled. Specifically, Eklund et al. (2016) demonstrated empirical FA rate control for a permutation test that involves the *two-sample* t-test. This is a nonparametric test for studies with a *between*-participants manipulation of the explanatory variable (e.g., patients-versus-controls, old-versus-young, treatment-versus-placebo). Different from a study with a within-participants manipulation, every participant is now observed in only a single condition. The reference distribution of this two-sample permutation test is obtained by randomly permuting the biological data over the different experimental conditions. Crucially, decisions based on the resulting permutation p-value control the empirical FA rate.

Thus, depending on the nonparametric test, the empirical FA rate is either controlled (the two-sample permutation test) or is not controlled (sign-flipping test). In this paper, we will present an alternative for the sign-flipping test, and will also give a formal proof of this alternative’s FA rate control. This proof is highly similar to a proof of the FA rate control of the two-sample permutation test, which we will also give. Both proofs demonstrate FA rate control under the null hypothesis of statistical independence between the biological data and the explanatory variable.

### A Randomization Test Alternative for the Sign-Flipping Test that Controls the FA Rate

We propose a randomization test for a study with a within-participants manipulation of the explanatory variable. This test has two essential ingredients: multiple condition orders that reflect the effect of interest, and (2) random assignment of participants to these condition orders. Starting with the first ingredient, the multiple condition orders are obtained from a partitioning of the event times in as many partitions as the number of experimental conditions. Every partition of event times is then assigned to one experimental condition. For example, in a study with 8 event times and 2 experimental conditions (A and B), the two partitions of event times could be {1,2,5,7} and {3,4,6,8}, and this would result in the following pair of condition orders: [AABBABAB, BBAABABA]. This scheme generalizes to studies with more than two conditions: with k experimental conditions, there are k partitions of the event times and k condition orders. For simplicity, in the following, we restrict ourselves to two conditions.

The FA rate of the randomization test is controlled for all pairs of condition orders. However, the precise null hypothesis that is tested does depend on the pair of condition orders. This is clear from the fact that the condition orders can be chosen such that possible confounds are either prevented or allowed for. In studies with a within-participants manipulation of the explanatory variable, both order and expectancy confounds can occur, but it is also easy to control for them. Specifically, the condition orders can be chosen such that (1) the conditions are uncorrelated with their order (e.g., one condition should not dominate the first or the second half of the experiment), and (2) the condition orders contain no obvious regularity that induces expectancy effects in the participant (e.g., alternating conditions).

The second essential ingredient of the randomization test is the random assignment of participants to the possible condition orders. This is similar to a randomized between-participants experiment in which the participants are randomly assigned to the conditions, whereas here they are randomly assigned to the possible condition *orders*. Typically, this random assignment involves calls to a pseudo-random number generator. This random assignment is essential, because the randomization mechanism that is used for the assignment will also be used for calculating the randomization p-value (see further).

We now list the remaining steps of the randomization test:

1. Collect the data. These data are depicted schematically in Figure 1. The figure is schematic and applies to both scalar and high-dimensional biological data; the structure of the data arrays is not shown in the figure. Note that there are only two possible condition orders, but that the participants can have different time courses of triangle heights. This corresponds to individual differences in the responses to the stimuli.
2. Separately for every participant, quantify the effect of the experimental conditions (A versus B). This has been described in the previous.
3. Calculate a test statistic. This could be the one-sample t-statistic that is used in the parametric framework, but it could also be the simple average of the effect quantifications, calculated over participants.
4. Calculate a randomization p-value for the observed test statistic. This randomization p-value is calculated under a reference distribution that is obtained by randomly reassigning the participants to one of the two condition orders, while keeping the observed biological data fixed. Thus, for calculating this reference distribution, we make use of the same randomization mechanism that was used for the initial (prior to the data collection) assignment of the participants to the condition orders. Every random re-assignment is then combined with the observed biological data, and the test statistic is recalculated as if this re-assignment were the initial assignment. By repeating these steps (random re-assignment and recalculation) a large number of times (in principle, an infinite number of times), we obtain the randomization distribution, the reference distribution of a randomization test.
5. If the randomization p-value is less than some nominal alpha level, reject the null hypothesis of statistical independence between the biological data and the explanatory variable (here, represented by the two condition orders).

**Figure 1:**
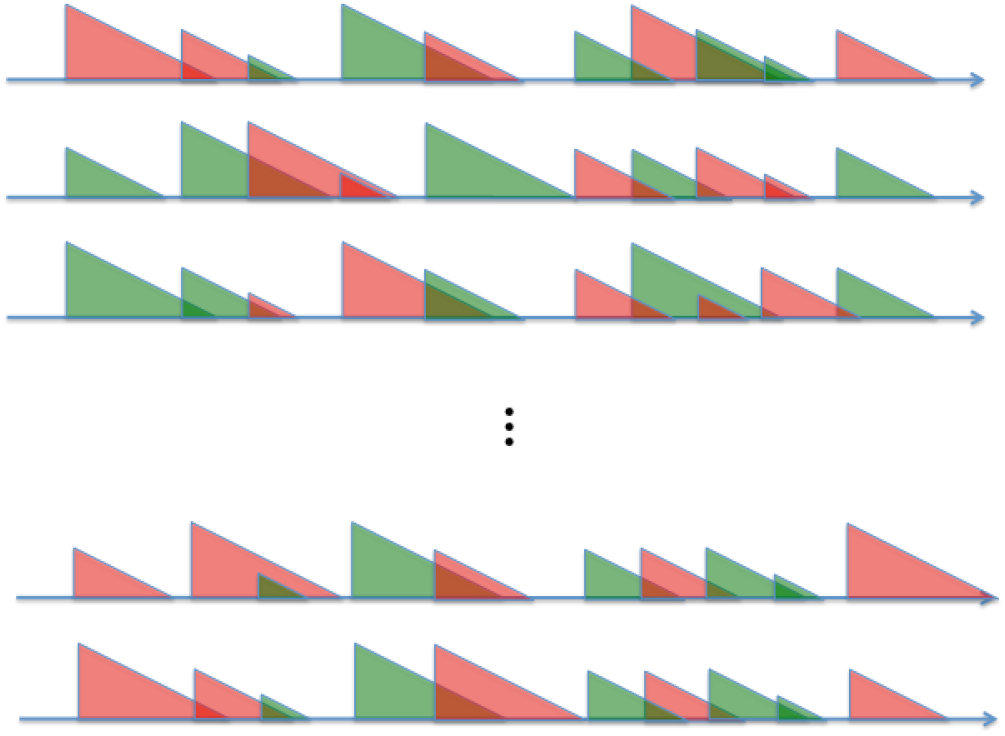
Schematic representation of the data of a study with a within-participants manipulation of the explanatory variable. Every timeline (row) corresponds to one participant and every triangle to one event. The colors of the triangles denote the experimental conditions (red=A, green=B), and their heights denote the amplitude of the biological data.

### A Comparison Between the Randomization and the Sign-Flipping Test

There are two important differences between the randomization and the sign-flipping test. First, the randomization test requires *two* condition orders whose contrast reflects the effect of interest. The sign-flipping test, on the other hand, can be applied both when a single or when multiple condition orders are used.

The second difference between the two tests is that the randomization test requires that the condition orders are randomly assigned to the participants, whereas the sign-flipping test can also be applied if they are assigned using a non-random mechanism. The random assignment requirement follows from the fact that statistical independence (our null hypothesis) is only defined between two random variables, and not between a random and a fixed (non-random) variable. Thus, the null hypothesis of statistical independence dictates random assignment, and this results in a randomization test.

In the following two sections, we will present the theoretical concepts that motivate the use of the randomization test. Specifically, we will give a proof of its FA rate control under the null hypothesis of statistical independence between the biological data and the explanatory variable. For the sake of clarity, we will first give this proof for studies in which the explanatory variable is manipulated *between* participants. In the section thereafter, we will show that this proof also applies to studies in which the explanatory variable is manipulated *within* participants. Finally, in the last section, we will demonstrate how a permutation test allows a test of the null hypothesis of statistical independence between the biological data and a *natural* explanatory variable.

## Randomization Tests for Studies with a *Between*-Units Manipulation of the Explanatory Variable

In this section, we will describe randomization tests that can be applied to both single- and multi-participant studies with a between-units manipulation of the explanatory variable. We make use of the concept of a *unit* because this allows for a general description that applies to both single- and multi-participant studies. In a single-participant study, the units are event times, and in a multi-participant study, they are participants.

When the units are event times, these times may indicate either brief stimulus presentations lasting less than a single scan, or the start of a longer stimulus presentation lasting for several scans. Such longer periods are often called “trials” or “blocks”. We will not distinguish between such trial-based studies and studies with single-scan events. The data of both study types are also analyzed in the same way: a GLM is fitted to the data, with the core regressors being stimulus functions convolved with a hemodynamic response function. Trial-based studies and studies with single-scan events only differ with respect to their stimulus functions: boxcars for trial-based, and a stick functions for studies with single-scan events.

When the units are participants, a between-units manipulation implies that all event times within a given participant are observed in the same experimental condition. This differs from studies in which the explanatory variable is manipulated *within* participants.

Different from the previous section, we will now introduce some notation. This allows us to provide a formal proof of the FA rate control of the randomization tests.

### Notation

The biological data is denoted by *Y*, and the explanatory variable by *X*. The variables *Y* and *X* are assumed to be random variables, which implies that they are the result of a random process. The values that were actually observed are called the random variables’ realizations, and they are denoted by, respectively, *y* and *x*.

One of the important facts demonstrated in this section, is that the FA rate control of a randomization test does not depend on any assumption about the probability distribution of *Y*. However, to keep the exposition concrete, it helps if we describe the different situations that correspond to the two unit types. For the first unit type (participants), it is easy to conceptualize a nonparametric test that involves random permutations of participant-specific component data structures. The latter is not possible for the second unit type (event times). However, our randomization test does not require such permutations of participant-specific component data structures, and therefore it applies to both unit types.

First, if the units are participants, then the variable *Y* is an array of *n* smaller statistically independent component data structures *Y*_*r*_ (*r* = 1, …, *n*), each one corresponding to a single participant. This situation is depicted in *Figure 2: Schematic representation of the data of a study with a between-units manipulation of the explanatory variable. Panel A depicts a study in which the units are participants, each of which is represented by one stick. Panel B depicts the data of a single-participant fMRI study in which the units are event times. Every event is represented by one triangle. The colors of the sticks and the triangles denote the experimental conditions and their heights denote the amplitude of the biological data*.

A, in which the lengths of the sticks correspond to the amplitudes of the components *Y*_*r*_, which typically are regression coefficients. The figure is schematic and applies to both scalar and high-dimensional biological data; the structure of the data arrays is not shown in the figure.

**Figure 2:**
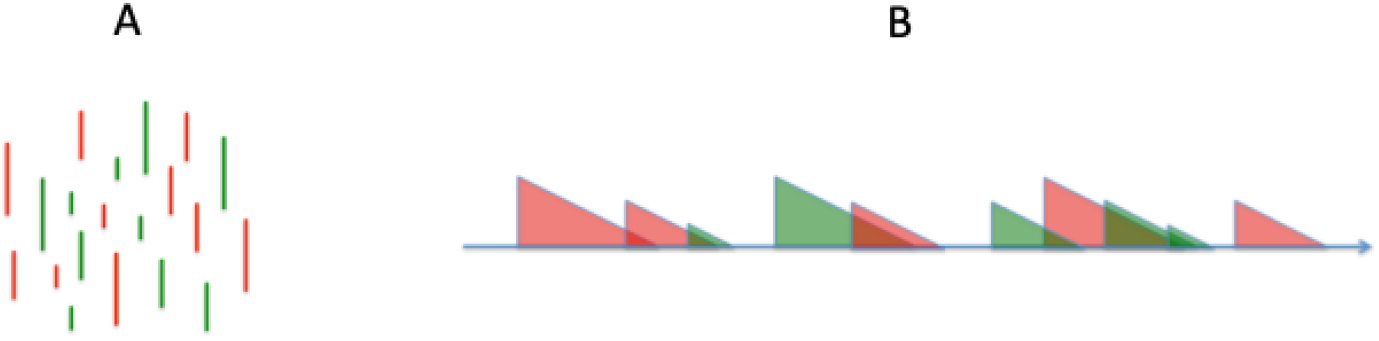
Schematic representation of the data of a study with a between-units manipulation of the explanatory variable. Panel A depicts a study in which the units are participants, each of which is represented by one stick. Panel B depicts the data of a single-participant fMRI study in which the units are event times. Every event is represented by one triangle. The colors of the sticks and the triangles denote the experimental conditions and their heights denote the amplitude of the biological data.

Second, if the units are event times in a single-participant fMRI study, it is not possible to separate *Y* in a set of *n* smaller statistically independent component data structures. This is because *Y* now is the whole MR time series obtained from that participant, with the effects of the different events superimposed on each other. This situation is depicted in Figure 2B, in which the heights of the triangles correspond to the magnitudes of the event-specific BOLD-responses.

The explanatory variable *X* is a variable of which the relation with the biological variable *Y* is of scientific interest: stimulus type, task/instruction, behavioral response, etc. In the context of a randomization test, the levels of this explanatory variable are experimental conditions to which the units are randomly assigned. The explanatory variable *X* is an array of *n* components *X*_*r*_ (*r* = 1, …, *n*), each one corresponding to a single unit. For example, in a single-participant study, *X*_*r*_ could indicate whether-or-not a predictive cue was given at the *r*-th event time. And in a multi-participant randomized experiment, *X*_*r*_ could indicate whether the *r*-th participant was assigned to the experimental or the control condition. In Figure 2, the realizations *x*_*r*_ of the component random variables *X*_*r*_ are depicted by the colors red and green.

### The Hypothesis of Statistical Independence

The randomization test we describe in this paper is a test of the hypothesis of statistical independence between the biological data *Y* and the explanatory variable *X*. This hypothesis pertains to the conditional probability distribution of the biological data *Y* given the explanatory variable *X*, which is denoted by *f*(*Y* = *y*|*X* = *x*), in which the lower-case letters *y* and *x* denote the realizations of the corresponding random variables. Now, if this conditional probability distribution depends on *x*, then we say that the explanatory variable has an effect on the biological variable. The probability distribution of *Y* may of course depend on other variables besides *X*, the explanatory variable of interest, but the effect of all these other variables will be considered noise that contributes to the variability of *Y* for a given realization *x* of *X*.

Formally, with a randomization test, we test the null hypothesis of statistical independence between *Y* and *X*:

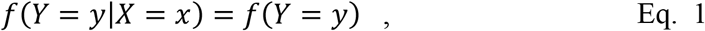

or, in brief, *f*(*Y*|*X*) = *f*(*Y*). In the remainder of this paper, unless there is a risk for confusion, we will disregard the distinction between a random variable (*X*, *Y*) and its realization (*x*, *y*). Statistical independence is symmetrical between *Y* and *X*, and therefore can also be expressed as follows:

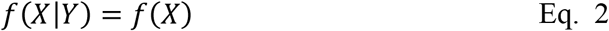

Eq. 2 is useful for proving the FA rate control of the randomization test.

Eq. 1 and Eq. 2 are the most general formulation of the null hypothesis of statistical independence between the biological data and the explanatory variable. More specific formulations are possible under suitable assumptions. For instance, under statistical independence between the unit-specific component data structures *Y*_*r*_, the null hypothesis can be formulated as follows:

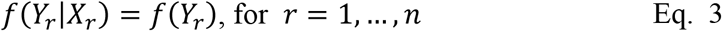

Here, the index *r* refers to the rank order of a unit that is randomly drawn from some population, and the functions *f*(*Y*_*r*_|*X*_*r*_) and *f*(*Y*_*r*_) therefore characterize probability distributions over this population. The formulation in Eq. 3 is useful when the units are participants. Instead, when the units are event time points in a single-participant fMRI study, no such component data structures can be specified, and only the general formulation in Eq. 1 and Eq. 2 is applicable.

It should be stressed that the variable *Y* and its components *Y*_*r*_ denote raw data. Of course, when calculating a test statistic, the raw data will be processed with the goal of extracting the relevant information for some phenomenon of interest (by means of averaging, GLM-based deconvolution, the Fourier transform, …). However, the theory of the randomization test has no implications for this data processing: if the null hypothesis of statistical independence holds for the raw data, it also holds for any function of the raw data. Of course, if this null hypothesis does *not* hold, then the choice of the test statistic may very well affect the probability of rejecting it (i.e., the sensitivity). In general, a well-informed choice of the test statistic (zooming in on the aspect of the data that is most likely to exhibit an effect) increases the sensitivity.

### Interpreting the Hypothesis of Statistical Independence

The null hypothesis *f*(*Y*|*X*) = *f*(*Y*) is rather abstract. However, the meaning of its ingredients *f*(*Y*|*X*) and *f*(*Y*) becomes more concrete by formulating them in terms of (1) the random processes that contribute to them (i.e., random sampling and within-participant randomness), and (2) one of the two units we are considering (i.e., participants and event time points). We start with discussing the contribution of random sampling.

In general, when units are randomly sampled from an infinite population, then repeating the study will result in a different sample with a different realization *y* of *Y*. The resulting probability distribution *f*(*Y*) therefore is characteristic of this population, and so is the null hypothesis of statistical independence, *f*(*Y*|*X*) = *f*(*Y*). When the units are participants that are randomly and independently sampled from some population, the equality *f*(*Y*_*r*_|*X*_*r*_) = *f*(*Y*_*r*_) indicates that, for the biological data of a randomly sampled participant, it does not matter in which experimental condition he/she is observed. Importantly, this null hypothesis pertains to a population of potential participants and not only to the observed. This contradicts the claim that a randomization test only makes an inference about the observed sample, and therefore would not allow one to generalize to a population (Nichols & Holmes, 2002, p. 8).

Strictly speaking, generalization to a well-defined population is only possible if the units are representative for that population. The common way to achieve this, is by sampling randomly from this population. However, in most neuroscience studies, participants are not randomly sampled from a well-defined population. Instead, most samples of participants are so-called convenience samples. This is a well-known problem that also affects parametric statistical inference, and therefore we will not consider it any further.

In a single-participant fMRI study, the units are event times within a participant. Because the effects of the different events are superimposed on each other, this study type does not involve unit-specific data components, and therefore can also not involve a physical process that samples these components randomly from some population. Given that the probability distributions *f*(*Y*) and *f*(*Y*|*X*) cannot result from random sampling, they must result from a source of randomness *within* the participant. This within-participant randomness refers to inherent randomness in the participant’s responses to the events: if the events would again be presented to the participant, different biological data would be observed, even with an identical realization of the explanatory variable.

Researchers typically formulate hypotheses in terms of the amplitudes of the stimulus-evoked hemodynamic responses (HRs). This differs from our null hypothesis *f*(*Y*|*X*) = *f*(*Y*), which is formulated at the level of the observed biological data. However, HRs (always assumed to be stimulus-evoked in the following) can be considered as random variables whose probability distribution may depend on the experimental conditions. Therefore, it is possible to formulate the null hypothesis as statistical independence between the random HRs and the explanatory variable. In the following subsection, we will show that, from this null hypothesis at the level of the random HRs, it follows that also the observed biological data *Y* and the explanatory variable *X* are statistically independent.

### From a null hypothesis at the level of HRs to a null hypothesis at the level of the whole recorded biological signal

We now introduce a small formal framework in which we express the observed biological data as a function of HR parameters (amplitude, delay, duration, …), event times, and experimental conditions. Researchers may be more interested in a null hypothesis with respect to HR parameters than a null hypothesis with respect to the whole recorded biological signal. We will use this formal framework to demonstrate that a null hypothesis with respect to the HR parameters implies a null hypothesis with respect to the whole recorded signal. Therefore, rejecting the latter null hypothesis implies that also the former must be rejected. We will use this framework for all study types: single- and multi-participant, involving between- and within-participant manipulations of the explanatory variable.

We start with a multi-participant study with a between-participant manipulation of the explanatory variable. We first introduce the event times as a vector *E*. For simplicity, we assume this vector to be identical for all participants, although it is easy to generalize our exposition to the case where *E* is random (but identically distributed in the two conditions). Next, we specify random HR parameters *C*_*r*_, jointly for all events. The random variable *C*_*r*_ pertains to a randomly drawn participant *r*. For a single voxel, one way to conceive *C*_*r*_ is as a matrix with dimensions parameter order and number of events. Possible parameters for single events are HR amplitude, delay and duration. We do not fully specify the HR parameters because there is no need for it, and leaving them underspecified highlights the generality of the theory. For instance, this underspecification allows to conceptually include parameter-depend ways of combining multiple event-specific HRs (possibly involving supra- or super-additive combination rules).

The biological data of a random participant depend on two signal components: a *combined HR* (CHR) and noise. A CHR is the outcome of a function *CHRF*(*E*, *C*_*r*_, *X*_*r*_) that takes as its input (1) the set of event times *E*, (2) the random HR parameters *C*_*r*_ for these event times, and (3) the explanatory variable *X*_*r*_. The functional form of this CHR does not have to be specified. The random biological data *Y*_*r*_ is generated as follows:

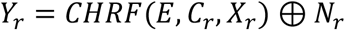

In this equation, *N*_*r*_ denotes noise, and ⊕ denotes an operator that combines the two signal components. This operator may denote simple addition, but also a supra- or super-additive combination. We assume that the noise *N*_*r*_ is independent of the explanatory variable *X*_*r*_:

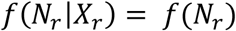

We want to test the null hypothesis that the HR parameters *C*_*r*_ are statistically independent of the conditions. Formally,

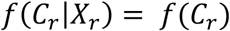

Because the probability distribution of *Y* depends on *X* only via *CHRF*(*E*, *C*_*r*_, *X*_*r*_), statistical independence at the level of the HR parameters *C*_*r*_ implies statistical independence at the level of the observed biological data *Y*. In other words, *f*(*C*_*r*_|*X*_*r*_) = *f*(*C*_*r*_) implies *f*(*Y*_*r*_|*X*_*r*_) = *f*(*Y*_*r*_). The latter hypothesis can be tested using the randomization test that is described in the following. Rejecting *f*(*Y*_*r*_|*X*_*r*_) = *f*(*Y*_*r*_) implies that also *f*(*C*_*r*_|*X*_*r*_) = *f*(*C*_*r*_) must be rejected.

Next, we consider a single-participant study with a between-event-times manipulation of the explanatory variable. Because we are no longer considering a participant that is randomly drawn from some population, in the formulae above, we must replace *Y*_*r*_, *X*_*r*_, *C*_*r*_ and *N*_*r*_ by, respectively, *Y*, *X*, *C* and *N*. Different from a multi-participant study, the event times *E* are now randomly partitioned into two sets, of which one is associated with condition A, and the other with condition B. This association of the event times with the experimental conditions is specified in the vector *X*. As an aside, every event time in a set could in principle be augmented with a different event duration, but this has no consequences for the properties of the statistical test. In the following, we will only consider the simple case of equal event durations.

With this change of the symbols, we are almost ready with reusing our explanation for a multi-participant study to single-participant one: rejecting *f*(*Y*|*X*) = *f*(*Y*) implies that also *f*(*C*|*X*) = *f*(*C*) must be rejected. It must be stressed that the probability distributions *f*(*Y*|*X*), *f*(*Y*), *f*(*C*|*X*), *f*(*C*), *f*(*N*(|*X*), and *f*(*N*) are all specific for the participant in this single-participant study; they all refer to probabilities under hypothetical replications of this single-participant study.

### The Randomization Test

A randomization test can be performed with an arbitrary test statistic *S*(*y*^*obs*^, *x*^*obs*^), in which *y*^*obs*^ and *x*^*obs*^ are the realizations of *Y* and *X* that were observed in the study. In a multi-participant study (in which the units are the participants), the test statistic is typically based on the difference between group averages of regression coefficients, with every group corresponding to one condition. In a single-participant study (in which the units are event times), the test statistic is typically based on a difference (contrast) between condition-specific regression coefficients obtained from a single-participant GLM-analysis.

The reference distribution for the test statistic is obtained by repeatedly calling the same randomization mechanism that also generated *x*^*obs*^, and plugging the resulting random variable *X* in the test statistic: *S*(*y*^*obs*^, *X*). There is a risk for confusion here, because we use the symbol *X* both to denote the random variable that generates the initial assignment *x*^*obs*^, as well as the random variable that is used to construct the reference distribution under which the p-value is calculated (using the fixed values *x*^*obs*^ and *y*^*obs*^). In the following, whenever there is a risk for confusion, we will use *X*^*rand*^ to denote the random variable that is used to construct the reference distribution. We will use the same name (randomization distribution) to denote *f*(*X*), *f*(*X*^*rand*^), and the reference distribution *f*(*S*(*y*^*obs*^, *X*^*rand*^)). The p-value is calculated by evaluating *S*(*y*^*obs*^, *x*^*obs*^) under *f*(*S*(*y*^*obs*^, *X*^*rand*^)).

The randomization distribution *f*(*X*) is determined by the researcher, and for the implementation of a randomization test, it is required to know the probabilities of each of the different random assignments. In terms of the elements of Figure 2, drawing from the randomization distribution means that the colors red and green are randomly reassigned over a fixed set of sticks or triangles. Most often, the researcher performs the random assignment by randomly drawing without replacement from a group of condition labels that contains the desired numbers of labels for each of the conditions (e.g., 10 As and 10 Bs). However, other randomization mechanisms, such as randomly drawing *with* replacement, are also possible, and our theory applies to all of them.

When randomly drawing without replacement from a group of *n* condition labels with an equal number for each of the two conditions, then every unique distribution of the conditions over the *n* units has a probability of 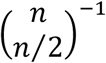. When *n* = 20 (Figure 2a), this probability equals 5.4*e* − 6, and when *n* = 10 (Figure 2b), it equals 4.0*e* − 3. When randomly drawing *with* replacement from the same group of condition labels, every unique distribution of the conditions over the *n* units has a probability of 0.5^*n*^. The difference between random draws with and without replacement is not relevant for the following.

To exactly construct the randomization distribution, all possible assignments must be enumerated. When the number of units is large, it is computationally infeasible to perform a complete enumeration. However, in this situation, it is possible to approximate the randomization distribution (with arbitrary accuracy) by randomly drawing values from it. The resulting approximation is denoted as a Monte Carlo estimate, and its accuracy can be quantified by means of a Monte Carlo confidence interval.

The decision about the null hypothesis is taken on the basis of a p-value that is obtained under the randomization distribution. For a test statistic of which large values provide evidence against the null hypothesis, the randomization p-value can be expressed as *P*(*S*(*y*^*obs*^, *X*) > *S*(*y*^*obs*^, *x*^*obs*^)), in which *P* denotes “probability”.

The decision about the null hypothesis (accept or reject) is taken by comparing the randomization p-value with the so-called *nominal alpha level*. This nominal alpha level is some a priori value between 0 and 1, typically 0.05 or 0.01. If the randomization p-value is less than the nominal alpha level, the null hypothesis is rejected; otherwise, it is accepted.

### The Randomization Test Controls the FA Rate

We will now prove that, under the null hypothesis of statistical independence between *X* and *Y*, the probability of a randomization test rejecting this null hypothesis is equal to the nominal alpha level. We will demonstrate this FA rate control in a different way as for a classical parametric statistical test (e.g., a t-statistic). In the latter case, the test statistic’s reference distribution (its probability distribution under the null hypothesis) is known prior to collecting the biological data. In contrast, for a randomization test, the reference distribution depends on *y*^*obs*^. We will deal with this dependence in two steps:

1. We start by proving FA rate control for a specific realization *y*^*obs*^ of Y. That is, we will prove *conditional* FA rate control.
2. We prove that conditional FA rate control implies unconditional FA rate control (i.e., independent of *y*^*obs*^).

#### The randomization test controls the FA rate conditionally given Y = y^obs^

The FA rate is the probability of falsely rejecting the null hypothesis. A false rejection occurs if, under this null hypothesis, the randomization p-value *P*(*S*(*y*^*obs*^, *X*) > *S*(*y*^*obs*^, *x*^*obs*^)) is less than the nominal alpha-level (*α*). The probability of a false rejection (i.e., the FA rate) is evaluated over hypothetical replications of the study, and therefore we must allow for the possibility that the initial assignment (explanatory variable) *x*^*obs*^ differs over these replications. We begin by fixing *Y* at *y*^*obs*^, and will therefore consider the conditional FA rate given *Y* = *y*^*obs*^. Now, the randomization p-value for a given study is *P*(*S*(*y*^*obs*^, *X*^*rand*^) > *S*(*y*^*obs*^, *X* = *x*^*obs*^)), in which the probability is taken over the realizations of *X*^*rand*^. For given values of *y*^*obs*^ and *x*^*obs*^, this p-value is a constant, but as a function of the random variable *X*, it is a random variable. Now, the probability of rejecting the null hypothesis equals the probability that this random p-value is less than *α*. In terms of the random test statistic *S*(*y*^*obs*^, *X*), this equals the probability that *S*(*y*^*obs*^, *X*) is larger than the (1 − *α*) × 100 percent quantile of the randomization distribution *f*(*S*(*y*^*obs*^, *X*^*rand*^)). Here, we tacitly assume a one-tailed test. However, our proof easily generalizes to two-tailed tests.

Because our objective is to determine the FA rate conditionally given *Y* = *y*^*obs*^, we must know the corresponding conditional probability distribution of *S*(*y*^*obs*^, *X*): *f*(*S*(*y*^*obs*^, *X*)|*Y* = *y*^*obs*^). At this point, we make use of the null hypothesis of statistical independence between *X* and *Y*. Specifically, because *S*(*y*^*obs*^, *X*) is a function of the random variable *X*, under this null hypothesis, also *S*(*y*^*obs*^, *X*) is statistically independent of *Y*. Thus, the conditional probability distribution *f*(*S*(*y*^*obs*^, *X*)|*Y* = *y*^*obs*^) is identical to *f*(*S*(*y*^*obs*^, *X*)), which in turn is identical to the randomization distribution *f*(*S*(*y*^*obs*^, *X*^*rand*^)), whose (1 −*α*) × 100 percent quantile is used to determine whether the null hypothesis will be rejected. As a consequence, under the null hypothesis, and conditional on *Y* = *y*^*obs*^, the probability that *S*(*y*^*obs*^, *X*) is larger than the (1 − *α*) × 100 percent quantile of the randomization distribution *f*(*S*(*y*^*obs*^, *X*^*rand*^)) is exactly equal to *α*. In other words, conditional on *Y* = *y*^*obs*^, the probability of falsely rejecting the null hypothesis is exactly equal to *α*. This completes our proof of the fact that the randomization controls the FA rate conditionally given *Y* = *y*^*obs*^.

The conditional probability distribution *f*(*Y*, *X*|*Y* = *y*^*obs*^) is depicted schematically in Figure 3, separately for participants (panel A) and for event times (panel B) as units. The black vertical bars denote the fact that we are considering a conditional distribution, and the grey sticks on the right-hand side of the bars depict *Y* = *y*^*obs*^, the random variable on which we condition. The colored sticks and triangles on the left-hand side depict (*y*^*obs*^, *X*), the argument of the test statistic. Crucially, the biological variable is fixed at its observed value *y*^*obs*^, and the explanatory variable *X* is random. Under the null hypothesis, *X* follows the randomization distribution *f*(*X*). This can be visualized by changing the colors as governed by *f*(*X*), while keeping the length of the sticks constant. Crucially, *f*(*X*) equals *f*(*X*^*rand*^), the distribution under which the p-value is calculated.

**Figure 3:**
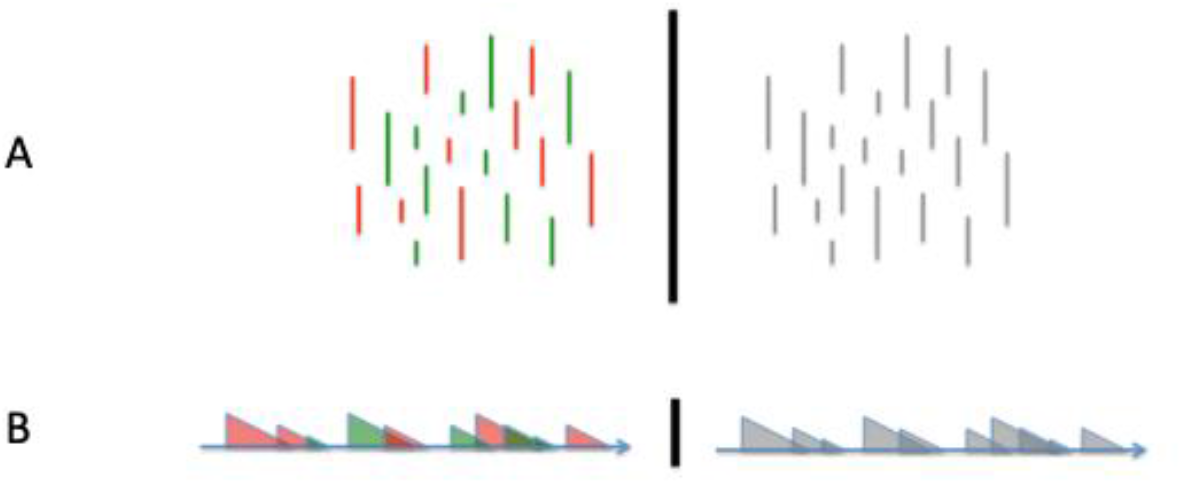
Schematic representation of the conditional probability distribution of a study with a between-units manipulation of the explanatory variable. Panel A depicts a study in which the units are participants, each of which is represented by one stick. Panel B depicts the data of a single-participant fMRI study in which the units are event times. Every event is represented by one triangle. The colors of the sticks and the triangles denote the experimental conditions and their heights denote the amplitude of the biological data. The grey sticks and triangles on the right-hand side of the vertical bars depict the biological data, which is the random variable on which we condition.

#### FA rate control conditionally given Y = y^obs^ implies unconditional FA rate control

At first sight, controlling the FA rate in this conditional sense (i.e., conditional on *Y* = *y*^*obs*^) is not very appealing. After all, who is interested in the conditional FA rate given one specific realization of *Y*? However, the FA rate is equal to the critical alpha-level, regardless of whether the p-value has a conditional or an unconditional interpretation. This is because, for every realization *y*^*obs*^ of *Y* on which we condition, the FA rate is equal to the same critical alpha-level. Therefore, if we average over the probability distribution of the random variable *Y*, the FA rate remains equal to this critical alpha-level. This can also be shown in a short derivation. In this derivation, the FA rate under the conditional distribution *f*(*Y*, *X*|*Y* = *y*^*obs*^) is denoted by *P*(Reject *H*_0_|*Y* = *y*^*obs*^), and the FA rate under *f*(*Y*, *X*) by *P*(Reject *H*_0_). The FA rate *P*(Reject *H*_0_) is obtained by averaging the conditional FA rate *P*(Reject *H*_0_|*Y* = *y*^*obs*^) over the probability distribution *f*(*Y* = *y*^*obs*^):

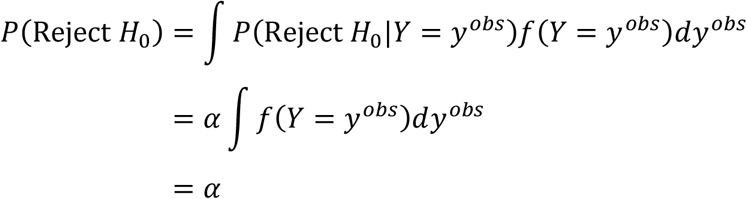

In the first line of this derivation, we make use of the following equality from elementary probability theory: *P*(*A*) = ∫ *P*(*A*|*B* = *b*)*P*(*B* = *b*)*db*. And in the third line, we make use of the fact that the probability densities *f*(*Y* = *y*^*obs*^) integrate to 1.

We can conclude that an FA rate that is controlled under the conditional distribution *f*(*Y*, *X*|*Y* = *y*^*obs*^) is also controlled under the corresponding unconditional distribution *f*(*Y*, *X*). This conclusion is a special case of the following general fact: for every event (in our case, falsely rejecting the null hypothesis) whose probability is controlled under a conditional distribution, also the probability under the corresponding unconditional distribution is controlled. This general fact will be called the “conditioning rationale”.

The conditioning rationale is used to prove the unconditional control of the FA or type 1 error rate, and does not involve a claim about the type 2 error rate (i.e., the probability that null hypothesis is maintained while in fact the alternative hypothesis is true). This is similar to classical parametric statistics, in which only the type 1 error rate is controlled. However, different from classical parametric statistics, in the nonparametric framework the researcher is free to choose the test statistic. He may do this on the basis of prior knowledge, with the objective to reduce the type 2 error rate.

## Randomization Tests for Studies with a *Within*-Participants Manipulation of the Explanatory Variable

We now consider studies with two levels of units (participants and event times) in which one level (event times) is observed within the other (participants). To investigate the effect of an explanatory variable, every participant is observed in all the experimental conditions (the different levels of the explanatory variable). This is achieved by assigning the within-participants units (event times) to different conditions.

### Notation and Assumptions

The biological variable *Y* is an array of *n* smaller component data structures *Y*_*r*_ (*r* = 1, …, *n*), each one corresponding to one participant that is randomly and independently drawn from some population. Similarly, *X* is an array of *n* components *X*_*r*_ (*r* = 1, …, *n*), each one corresponding to one participant. Every component *X*_*r*_ in turn consists of *m* subcomponents *X*_*rs*_ (*s* = 1, …, *m*), each one corresponding to one event time. In the following, we will denote *X*_*r*_ as the *condition order*.

### The Hypothesis of Statistical Independence

We test the hypothesis of statistical independence between the explanatory variable *X* and the biological data *Y*: *f*(*X*|*Y*) = *f*(*X*). Because the *n* participants are randomly and independently drawn from some population, the null hypothesis of statistical independence also holds at the level of a randomly drawn participant (indexed by *r*): *f*(*X*_*r*_|*Y*_*r*_) = *f*(*X*_*r*_). In words, the biological data *Y*_*r*_ of a randomly drawn participant do not inform us about his condition order *X*_*r*_.

This null hypothesis depends on the randomization distribution *f*(*X*_*r*_). Specifically, the randomization distribution determines the generality of the inference that follows from the rejection of the null hypothesis of statistical independence. Here, “generality” refers to the population from which the participants are randomly drawn. As will be explained in a later section (*Selecting a Randomization Distribution*), the smaller the number of possible realizations of the random variable *X*_*r*_, the more general the inference that follows from the rejection of the null hypothesis.

### The Randomization Test and its FA Rate Control

The randomization test for a study with a within-participants manipulation of the explanatory variable is very similar to the one for studies with a between-units manipulation. To construct *f*(*S*(*y*^*obs*^, *X*^*rand*^)), the randomization distribution of the test statistic, we separately and independently draw from the participant-specific randomization distribution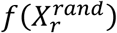, which generates the rows of the participant-by-unit matrix *X*^*rand*^.

The proof of the randomization test’s FA rate control is identical to the proof that was given in the context of a study with between-units manipulations of the explanatory variable. This proof is also applicable here because it does not depend on any assumption about the probability distribution of *Y*, nor an assumption about the structure of *X*_*r*_ (i.c., whether it is a scalar or a vector).

However, the randomization tests for the two study types (with a between-units and a within-participants manipulation of the explanatory variable) differ with respect to the test statistic *S*(*y*^*obs*^, *x*^*obs*^). In a study with a within-participants manipulation, the test statistic is typically based on the contrasts between condition-specific regression coefficients that are obtained from the multiple participant-specific GLM-analyses. These contrasts are combined over the participants, by averaging or by calculating a one-sample (paired-samples) t-statistic.

### Selecting a Randomization Distribution *f*(*X*_*r*_)

The randomization distribution *f*(*X*_*r*_) determines the generality of the inference that follows from the rejection of the null hypothesis of statistical independence. The concept of generality of inference is related to the distinction between fixed and random effect hypotheses in parametric statistics. Specifically, rejecting a random effect null hypothesis corresponds to a general inference, and rejecting a fixed effect one corresponds to a specific inference. In the following, we first prime the proper intuition by describing the parametric fixed and random effect statistical tests. Second, we give a more detailed description of the concept of generality of inference. Third, we describe how to deal with sensitivity and confounds when selecting possible condition orders. Fourth and last, we motivate a particular randomization distribution (involving a pair of complementary condition orders) using our simple formal framework in which we express the observed biological data as a function of HR parameters, event times, and experimental conditions.

#### Parametric fixed and random effect statistical tests

Although there are no universally accepted definitions of fixed and random effects (Gelman, 2005), in neuroimaging one typically follows the definitions in the publications by members of the Wellcome Center for Human Neuroimaging (e.g., Holmes & Friston, 1998). Following these definitions, we first describe a fixed effect statistical test for a study with two experimental conditions (A and B). The interest is in two-sided differences between the regression coefficients for these two conditions (A>B or B>A). There are multiple ways in which such a parametric fixed effect test can be performed, and here we describe one:

1. For every participant, fit a GLM with core regressors corresponding to the two experimental conditions. From the output of this analysis, calculate a t-statistic by dividing the regression coefficient contrast A-versus-B by its standard error.
2. Combine the t-statistics over the participants. There are two main ways of doing this. The first and most obvious way is by averaging the t-statistics. This combination has the disadvantage that, if there are between-condition differences in both directions (A>B and B>A), cancellation of positive and negative differences reduces sensitivity. The second way of combining the t-statistics, is by averaging the squared instead of the original t-statistics. In this way, cancellation of positive and negative differences is prevented.
3. Evaluate the combined test statistic under the reference distribution that is obtained under the combined null hypothesis of equal expected condition-specific regression coefficients for every individual participant. Under this null hypothesis, the average t-statistic has an asymptotic standard normal distribution, and the average of the squared t-statistics has an asymptotic scaled chi-square distribution (with degrees of freedom equal to the number of participants).

There are two ways to interpret the outcome of this fixed effect test, depending on whether or not the participants were randomly drawn from some population. If the researcher has selected just this specific set of participants (no random sampling), then the null hypothesis of this fixed effect test is actually a fixed set of participant-specific null hypotheses: equal expected condition-specific regression coefficients for every participant in the set. The expected values in these participant-specific null hypotheses are with respect to a probability distribution over independent replications of the same study with the same participants.

On the other hand, if the participants are randomly drawn from some population, then the outcome of this so-called fixed effect test is also relevant for this population. Specifically, the null hypothesis of this test now entails that the probability is *zero* that a randomly drawn participant has equal expected condition-specific regression coefficients (again with the expectation with respect to a probability distribution over independent replications with the same participant). This null hypothesis has been called the global null hypothesis by (Nichols, Brett, Andersson, Wager, & Poline, 2005). Clearly, although this null hypothesis pertains to a population of potential participants, it is very restrictive because it allows us only to infer that there are potential participants for which there is an effect, but not that this effect (e.g., A>B) generalizes to the population. This is most clear in studies with a large number of events, in which a participant-specific t-statistic will become very large if that participant has unequal expected condition-specific regression coefficients. Therefore, a single participant may dominate the outcome of the test. Moreover, even a single-participant (*n* = 1) study may result in a rejection of this restrictive population-level null hypothesis. This is an example of a very specific inference: a small subpopulation of participants with unequal expected condition-specific regression coefficients is sufficient to reject the null hypothesis.

We now consider a parametric random effect statistical test. This test is performed as follows:

1. For every participant, fit a GLM to the biological data, and calculate the regression coefficient contrast A-versus-B.
2. Calculate a one-sample (paired-samples) t-statistic on these participant-specific regression coefficient contrasts.
3. Evaluate the t-statistic under the reference distribution that is obtained under the assumption that the expected value of the participant-specific regression coefficient contrasts is equal to zero. Crucially, the expectation is now taken with respect to a probability distribution over some population of participants (instead of independent replications with the same participants). The reference distribution for this t-statistic is a Student T-distribution with number of degrees of freedom equal to the number of participants minus one.

The null hypothesis of this random effect test pertains to the expected value of the regression coefficient contrast, calculated over the population of potential participants. If this random effect null hypothesis is false, then so is the fixed effect null hypothesis (pertaining to the probability that a randomly drawn participant has equal expected values in his condition-specific populations of trials), but not the other way around. Specifically, the population of potential participants may contain two subpopulations, of which one has an effect in one direction (say, A>B) and the other has an effect in the other (B>A). The sizes of these subpopulations and their effect sizes could be such that there is a perfect cancellation of the effects, resulting in the random effect null hypothesis holding for the whole population.

This is an example of a null hypothesis that allows for a general inference. This is because it pertains to the whole population of potential participants. Specifically, rejecting this random effect null hypothesis implies that, on average over the potential participants, the regression coefficient contrast differs from zero in a particular direction.

#### The generality of inference allowed for by a null hypothesis of statistical independence

In the following, we will consider two different randomization distributions *f*(*X*_*r*_) for a study with a within-participants manipulation of the explanatory variable. These two randomization distributions correspond to null hypotheses that allow for inferences with different levels of generality. The first randomization distribution involves reusing the randomization distribution for a single-participant study with a between-event-times manipulation of the explanatory variable. In that study type, the researcher randomly assigns the event times to the conditions by randomly drawing (with or without replacement) from a group of condition labels. An important feature of that study type is the very large number of different condition orders. This number depends on the number of event times and the number of labels for each of the two conditions. With realistic values for the number of event times and the number of labels (a few hundreds), the number of different condition orders can become huge (e.g., 9.0549e+58 for 200 event times that are evenly split between two conditions).

Crucially, a large number of different realizations of the condition order *X*_*r*_ creates the opportunity for one or a few participants dominating the outcome of the randomization test. Specifically, if a single participant’s biological data 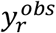 is strongly associated with the condition order 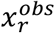, this results in a regression coefficient contrast that is much more extreme than what is expected under statistical independence (condition orders drawn from the randomization distribution). The contribution of this participant’s data to the group-level test statistic may also make this group-level test statistic very unlikely. This resembles evaluating the statistical significance of the average of 20 numbers: 19 numbers around 0 and one equal to 10. If these numbers’ probability distributions under the null hypothesis all have an expected value of 0, and a standard deviation that is much less than the extreme value 10 (corresponding to one participant-specific test statistic in the tail area of its randomization distribution), then also the average of these 20 numbers (0.5, which we take as our group-level test statistic) is unlikely under the null hypothesis that the expected value equals 0. Specifically, if the common standard deviation of these 20 null distributions is 1, then the standard deviation of this average is equal to 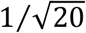, and the value 0.5 is in the 5% two-sided tails of its normal distribution. This is an example of a very specific inference because, for a rejection of the null hypothesis, it is sufficient that only one or a few participants have biological data that are statistically *de*pendent of the explanatory variable. This corresponds to a fixed-effects null hypothesis.

Next, we consider a randomization distribution *f*(*X*_*r*_) that generates only two possible condition orders. As will be shown, this randomization distribution allows for a general inference, which corresponds to a random-effects null hypothesis. For statistical sensitivity, the two condition orders must be chosen such that their contrast optimally reflects our effect of interest: the difference between the conditions A and B. The relevant information in the biological data *Y*_*r*_ pertains to the data’s association (over the events) with these conditions. Thus, for the appropriate sensitivity of the statistical test, we need a pair of condition orders that is maximally informative about this association. We achieve this by selecting a pair of condition orders of which the members are each other’s complement (e.g., AABBABAB and BBAABABA).

With only two possible condition orders, the null hypothesis *f*(*X*_*r*_|*Y*_*r*_) = *f*(*X*_*r*_) involves that the biological data *Y*_*r*_ are uninformative about whether the condition order is AABBABAB or BBAABABA. Finding evidence against this null hypothesis is more difficult than in a study with a large number of condition orders. With only two condition orders, finding sufficient evidence against the null hypothesis requires consistency among the participants. Specifically, if a single participant’s biological data 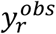 is strongly associated with the condition order 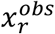, this cannot result in a regression coefficient contrast that is extreme under the null hypothesis. This is because the randomization distribution for a single participant has only two values, each one associated with one condition order. Therefore, finding sufficient evidence against the null hypothesis requires that the data from multiple participants must exhibit consistency (via their contribution to the test statistic) in the way they differentiate between the two condition orders (e.g., either A>B or B>A). Although the number of events per participant affects the reliability of the evidence for every individual participant, this does not reduce the need to combine this evidence over multiple participants.

#### Dealing with sensitivity and confounds when selecting possible condition orders

As explained in the above, the pair of condition orders is selected such that the statistical test has the appropriate sensitivity for detecting the effect of interest. However, the sensitivity is not only determined by whether or not the condition orders in a pair are complementary; it is also determined by the number of events and the inter-event times. With respect to the latter, the sensitivity depends on the correlation between the GLM regressors for the different conditions (stimulus functions convolved with a hemodynamic response function): if this correlation decreases from 1 to −1, then the sampling (noise) variance of the GLM regression coefficient contrast decreases from infinity to 0. Therefore, the inter-event times should be chosen such that the correlation between these GLM regressors is as small (negative) as possible.

Selecting a pair of condition orders of which the members are each other’s complement does not guarantee that all possible confounds are prevented. Possible confounds are order and expectancy effects. However, it is easy to control for these effects. Specifically, the condition orders can be chosen such that (1) the conditions are uncorrelated with their order (i.e., one condition should not dominate the first or the second half of the experiment), and (2) the condition orders contain no obvious regularity that induces expectancy effects in the participant (e.g., alternating conditions). Block randomization is an excellent method to generate a pair of condition orders that fulfills these requirements. This method starts by separating events in blocks (of size *k*) according to their temporal order: the first *k* events in block 1, the next *k* events in block 2, etc. Then, within every block, *k*/2 events are randomly assigned to one condition, and the remaining *k*/2 to the other (i.e., random sampling without replacement). With two blocks of size 4 (*k* = 4), the result could be AABBABAB. The second condition order is then taken as the complement of the first (i.e., BBAABABA).

In sum, we have introduced randomization distributions that correspond to null hypotheses that allow for inferences with different levels of generality; the larger the number of possible realizations (condition orders), the less general the inference that follows from a rejection of the null hypothesis. In addition, to achieve the appropriate sensitivity for detecting the effect of interest and to prevent confounds, care must be taken in selecting the possible condition orders.

As for a single-participant study, also for a multi-participant study, it is useful to consider the null hypothesis *f*(*X*_*r*_|*Y*_*r*_) = *f*(*X*_*r*_) as the result of a null hypothesis that is formulated at the level of the hemodynamic responses (HRs). In the following subsection, we demonstrate this for a randomization distribution with two possible realizations that are each other’s complement.

### From a null hypothesis at the level of HRs to a null hypothesis at the level of the whole recorded biological signal

We consider a random participant *r* with biological data *Y*_*r*_ and explanatory variable *X*_*r*_. The event times *E* are partitioned in two sets, *E*_1_ and *E*_2_, of which one is randomly associated with condition A, and the other with condition B. Our theory does not require that the event times *E* and/or their partitions *E*_1_ and *E*_2_ are identical for every participant. However, when the events (their times and their values) are under experimental control, there is no reason for different participants to have different event times *E* and/or partitions *E*_1_ and *E*_2_. The situation is different in studies in which the event times and/or their values are (partially) controlled by the participant (e.g., in a self-paced experiment and/or when the explanatory variable depends on behavior). We will return to this when we discuss natural explanatory variables (e.g., accuracy, reaction time, pupil diameter) and how their association with the biological data can be evaluated by means of permutation tests.

We specify random HR parameters *C*(*m*_1_) and *C*(*m*_2_), in which *m*_1_ and *m*_2_ are the number of events in, respectively, *E*_1_ or *E*_2_. As in a single-participant study, *C*(*m*_1_) and *C*(*m*_2_) can be conceived as matrices with dimensions *Parameter Order* and *Number of Events*.

We want to test the null hypothesis that the HR parameters *C*(*m*_1_) and *C*(*m*_2_) are statistically independent of the conditions. To express this formally, we introduce the scalar random variable *Z*, which indicates the experimental conditions A and B (thus, *Z* = *A* or *Z* = *B*). Now, the null hypothesis of interest can be expressed as follows:

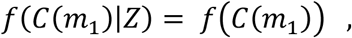

and

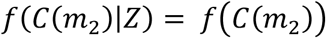

Crucially, and different from a single-participant study, the probability distributions *f*(*C*(*m*_1_)|*Z*) and *f*(*C*(*m*_2_)|*Z*) pertain to the population from which the participants were randomly sampled. The realizations of the random variables *C*(*m*_1_) and *C*(*m*_2_) exhibit variability in two levels: (1) across random draws from the population of participants, and (2) across events within a participant (i.e., the columns of the matrices *C*(*m*_1_) and *C*(*m*_2_)). This differs from a single-participant study, in which the corresponding probability distribution is *f*(*C*|*X*), in which *C* has one column for every event, each one corresponding to one element of the vector *X*. In this single-participant study, the realizations of the random variable *C* only exhibit variability across the within-participant events.

The biological data for a random participant (*Y*_*r*_) is the result of a combination of three components: two condition-specific *combined HRs* (CHRs) and noise. A combined HR is the outcome of a function that takes as its input (1) a set of event times, (2) random HR parameters for these event times, and (3) the condition identity. The CHR for events *E*_1_ assigned to condition A is denoted by *CHRF*(*E*_1_, *C*(*m*_1_), *A*), and the one for events *E*_2_ assigned to condition B is denoted by *CHRF*(*E*_2_, *C*(*m*_2_), *B*). The random biological data *Y*_*r*_ for this assignment are generated as follows:

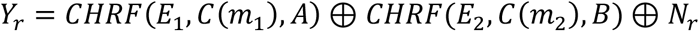

And for the reverse assignment, they are generated as follows:

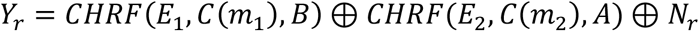

In these equations, *N*_*r*_ denotes noise, and ⊕ denotes an operator that combines the three signal components. This operator may denote simple addition, but also a supra- or super-additive combination.

We assume that the noise *N*_*r*_ is independent of the conditions:

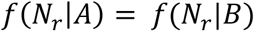

We also assume the operator ⊕ to be *commutative*: *Q* ⊕ *R* = *R* ⊕ *Q*.

For a random participant with event set *E*_1_ assigned to condition A, and event set *E*_2_ to condition B, we draw one realization from *f*(*C*(*m*_1_)|*Z* = *A*) and one from *f*(*C*(*m*_2_)|*Z* = *B*). Under the null hypothesis,

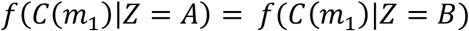

and

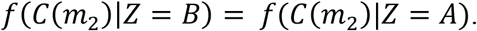

Therefore, under the null hypothesis, the random biological data *Y*_*r*_ for this assignment can also be generated as follows:

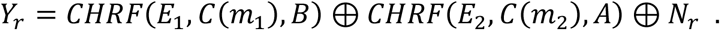

This is identical to the way *Y*_*r*_ is generated under the reverse assignment. Thus, from the null hypothesis of statistical independence at the level of the HR parameters, it follows that *f*(*Y*_*r*_|*X*_*r*_) = *f*(*Y*_*r*_), the null hypothesis at the level of the observed biological data. The latter null hypothesis can be tested using a randomization test, and if it is rejected, then so is the null hypothesis at the level of the HR parameters.

## Permutation Tests for Studies with a Natural Explanatory Variable

### Experimental versus Natural Explanatory Variables

An explanatory variable is called “experimental” if the researcher controls the assignment of the units to the levels of this variable. And it is called “natural” if it is not under experimental control. Only experimental variables can be randomized. Studies involving a natural explanatory variable are called “observational studies”.

We must distinguish between observational studies with a between-units and a within-participants manipulation of the explanatory variable. (Although a natural variable cannot be manipulated, for consistency, we will continue to use the term “manipulation” for all mechanism that controls this natural variable.) In a multi-participant study with a between-participants manipulation of the explanatory variable, possible natural explanatory variables are gender, age, disease status, education, ethnicity, and religion. And in a single-participant study with a between-event-times manipulation, possible natural explanatory variables are response accuracy, response time, and psychophysiological variables like pupil diameter. These same natural explanatory variables can also appear in a multi-participant study with a within-participants manipulation.

When a study involves a natural explanatory variable, the goal typically is to characterize the relation with a biological variable (Is there a relation, and what is its nature?), and not to evaluate whether there is a causal effect of the explanatory variable. In fact, the relation between the natural explanatory and the biological variable may very well be due to a third (confounding) variable. Here lies the strength of experimental explanatory variables: by randomly assigning the units to the experimental conditions, one ensures that there is no systematic association between pre-experimental characteristics of the units and the levels of the explanatory variable (Rubin, 1974). In other words, random assignment prevents confounding variables from creating a spurious association between the explanatory and the biological variable, and therefore allows for causal inference.

Experimental variables are almost always categorical (nominal), whereas natural variables can also be quantitative. For observational studies with a between-participants manipulation, familiar quantitative variables are age and performance level, and for observational studies with a between-event-times or a within-participants manipulation, they are response time and pupil diameter. For both types of observational studies (with a between-units and a within-participants manipulation of the explanatory variable), exactly the same permutation test can be used for categorical and quantitative variables.

In the next subsection, we present permutation tests that can be used to test the null hypothesis of statistical independence between a biological variable *Y* and a natural explanatory variable *X*. The main challenge in designing these permutation tests and in proving their FA rate control lies in the fact that the probability distribution of the natural explanatory variable *X* is unknown. We deal with this problem by drawing from a known *conditional* probability distribution of *X*. Importantly, this conditional probability distribution is only known under an assumption, and this assumption is different for observational studies with a between-units and those with a within-participants manipulation of the natural explanatory variable. The permutation tests for these two types of observational studies will be described in different subsections.

### Permutation Tests for Observational Studies with a Between-Units Manipulation of the Explanatory Variable

#### The permutation versus the randomization test

The permutation test is performed just like a randomization test, but now with a p-value that is calculated under the permutation distribution instead of the randomization distribution. This permutation distribution is obtained by randomly permuting the 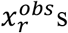 over the 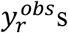. In terms of the elements of Figure 3a and 3b, this means that the fixed set of colors red and green are randomly permuted over, respectively, the sticks and the triangles. Every random permutation has the same probability equal to 1 over the total number of possible permutations. For our example in Figure 3a, this probability is 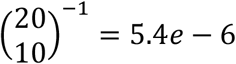. These probabilities in principle could also have been produced by a randomization mechanism, namely if the researcher had determined in advance to assign exactly 10 units to each of the two conditions. However, the crucial difference between a randomization and a permutation test is that, for the latter, the number of units in each of the conditions is not set in advance by the researcher, but observed from the units that were drawn. These numbers could also have been 9-vs-11, 8-vs-12, 7-vs-13, etc.

#### The permutation test controls the FA rate

We will now prove that a permutation test controls the FA rate at the nominal alpha level. This proof is along the same lines as the corresponding proof for a randomization test. Different from a randomization test, it is not required to know the probability distribution *f*(*X*). However, to prove the permutation test’s FA rate control, we must assume that the unit-specific explanatory variables *X*_*r*_ (*r* = 1, …, *n*) are statistically independent and identically distributed (*iid*). Crucially, under this assumption, the permutation distribution is the conditional probability distribution *f*(*X*|{*X*} = {*x*^*obs*^}): the probability of the ordered (*X*, the part before |) given the unordered components of *x*^*obs*^ ({*X*} = {*x*^*obs*^}, the part after |). Keep in mind that *X* and *x*^*obs*^ are arrays with components *X*_*r*_ and 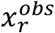, respectively.

##### 1. *f*(*X*|{*X*} = {*x*^*obs*^}) is a Permutation Distribution

We will now show that, under the assumption of *iid* components *X*_*r*_, *f*(*X*|{*X*} = {*x*^*obs*^}) is a permutation distribution. Under the assumption of *iid* components *X*_*r*_, *f*(*X*) can be written as follows:

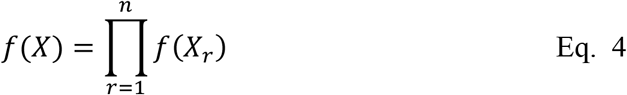

From Eq. 4, it follows that *f*(*X*) is *exchangeable*. Exchangeability means that the probability of explanatory variable *X* is invariant under permutation of the realizations *x*_*r*_ of the unit-specific random variables *X*_*r*_. Under exchangeability, *f*(*X*|{*X*} = {*x*^*obs*^}) is the permutation distribution. To see this in an example, consider a study with four units, such that 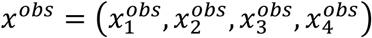. We assume that these four realizations are all different 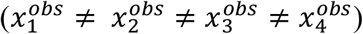, which is typically the case when the explanatory variable is quantitative. For this example, the unordered set {*x*^*obs*^} is the collection of all permutations of 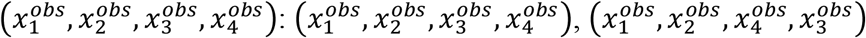, 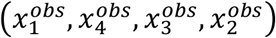, plus 21 more. Thus, the number of permutations is 4! = 24, and *f*(*X*|{*X*} = {*x*^*obs*^}) = 1/24, for each of the permutations of *x*^*obs*^. In many studies, some realizations 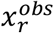 are equal (i.e., there are ties between the 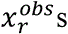), and this is typically the case when the explanatory variable is categorical. Ties reduce the size of the unordered set {*x*^*obs*^}. For example, if the interest is in gender differences, and the sample consists of two males and two females, then the number of unique components in the unordered set {*x*^*obs*^} is 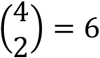. Consequently, *f*(*X*|{*X*} = {*x*^*obs*^}) = 1/6, for each of the unique permutations of *x*^*obs*^.

The reason for using *f*(*X*|{*X*} = {*x*^*obs*^}) instead of *f*(*X*) is that, contrary to the latter, *f*(*X*|{*X*} = {*x*^*obs*^}) is known. In the following, we use *L* to denote the number of unique values in the vector *x*^*obs*^. Then, under the assumption of statistically independent and identically distributed observations, *f*(*X*) can be written as follows:

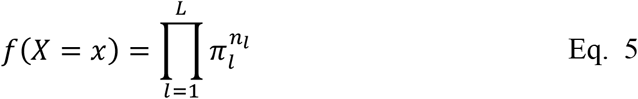

In Eq. 5, *π*_*l*_ is the probability of observing a unit in the *l*-th level (*l* = 1, …, *L*) of the explanatory variable, and *n*_*l*_ is the number of observations in this level. Now, when *X* is a natural explanatory variable, *f*(*X*) is unknown because the probabilities *π*_*l*_ are unknown. However, we now make use of the fact that the unordered observations {*X*} are a *sufficient statistic* for the unknown probabilities *π*_*l*_. A sufficient statistic is a function of a random variable (here, *X*) with the property that, by conditioning on its value, the resulting conditional probability distribution of the random variable becomes independent of the unknown parameters. To show that {*X*} is a sufficient statistic, we start from the fact that {*x*^*obs*^} (the observed realization of {*X*}) is equivalent to (*n*_1_, *n*_2_, …, *n*_*L*_), the number of observations in each of the *L* levels. Therefore, *f*({*X*} = {*x*^*obs*^}) is a multinomial distribution:

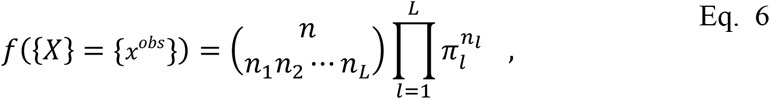

in which the first term on the right-hand side of Eq. 6 is the multinomial coefficient. Inserting Eq. 5 and Eq. 6 in the definition of a conditional probability, we obtain

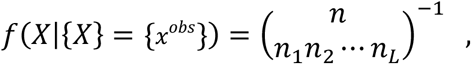

which is independent of the probabilities *π*_*l*_. Depending on the array (*n*_1_, *n*_2_, …, *n*_*L*_), there are several common special cases of the multinomial coefficient. For instance, when comparing two conditions, *L* = 2, and the multinomial becomes the binomial coefficient 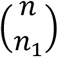 (remember that *n* = *n*_1_ + *n*_2_). Further, if every unit brings its own unique value 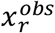 (which may happen if the explanatory variable is quantitative), the multinomial coefficient equals *n*!.

##### 2. FA Rate Control Using *f*(*X*|{*X*} = {*x*^*obs*^})

We can now complete our proof of the FA rate control of the permutation test, and we can do this along the same lines as the corresponding proof for the randomization test. The latter proof involved two steps, and this will also be the case for the current proof. In fact, the only difference between the two proofs is that, for the permutation test, we not only condition on *Y* = *y*^*obs*^ (as in the proof for the randomization test), but also on {*X*} = {*x*^*obs*^}. Specifically, our proof for the permutation test involves the following two steps:

1. The permutation test controls the FA rate conditionally given *Y* = *y*^*obs*^ and {*X*} = {*x*^*obs*^}.
2. FA rate control conditionally given *Y* = *y*^*obs*^ and {*X*} = {*x*^*obs*^} implies unconditional FA rate control.

Because our proof for the randomization test did not involve any component that prevented the conditioning on {*X*} = {*x*^*obs*^}, the same line of argument also provides the proof of the FA rate control for the permutation test.

The difference between a randomization and a permutation test can also be depicted schematically, and we did this in Figure 4. The right-hand side of the black vertical bar depicts *Y* = *y*^*obs*^ (the grey sticks) and {*X*} = {*x*^*obs*^} (the box with 10 red and 10 green balls). The numbers of red and green balls on the right-hand side could have been different (e.g., 14 red and 6 green balls), but their sum would always equal the number of sticks on the left- and the right-hand side. As a whole, this figure depicts the condition probability distribution *f*(*X*|*Y* = *y*^*obs*^, {*X*} = {*x*^*obs*^}), which is a permutation distribution. Without the box on the right-hand side, it would have been the randomization distribution *f*(*X*|*Y* = *y*^*obs*^). Drawing from the permutation distribution can be visualized by randomly assigning the colors in the box to the grey sticks.

**Figure 4:**
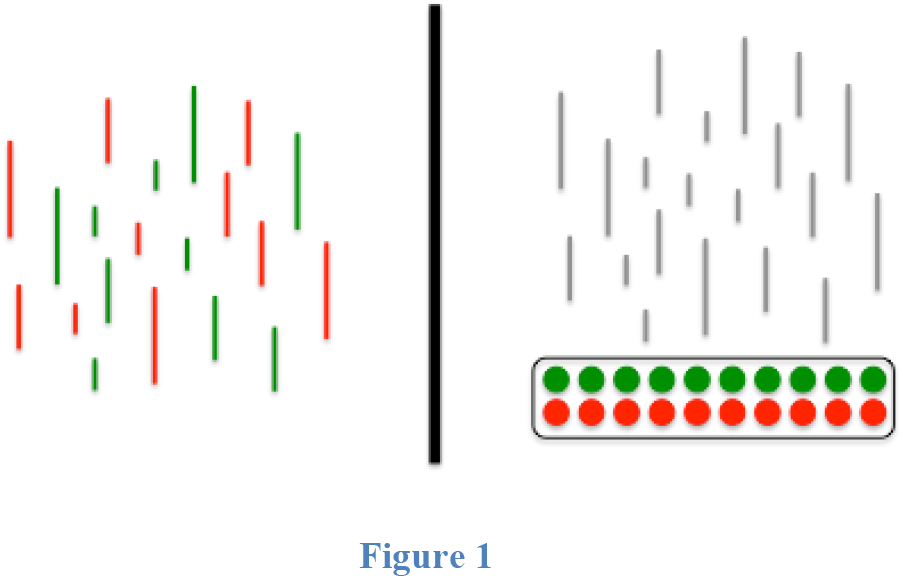
Schematic representation of the permutation distribution; the conditional probability distribution that is involved in a permutation test. The right-hand side of the black vertical bar depicts the biological data (the grey sticks) and the unordered components of the explanatory variable (the box with 10 red and 10 green balls).

### Permutation Tests for Observational Studies with a Within-Participants Manipulation of the Explanatory Variable

We now consider multi-participant observational studies in which the natural explanatory variable varies across the event times. Examples of such natural explanatory variables are response accuracy, response time, movement direction and psychophysiological variables like pupil diameter. As before, the probability distribution of the explanatory variable for a random participant *r* will be denoted by *f*(*X*_*r*_). This probability distribution is unknown and so are the possible realizations of *X*_*r*_. In fact, in an observational study, the number of events, the event times, and the associated conditions, can all be different for the different participants. For example, in a self-paced experiment in which accuracy will be the explanatory variable (with 0 and 1 denoting, resp., a correct and an incorrect response), the values of the explanatory variables for two participants could be [0,0,1,1,0] and [1,0,0,1,0,1]. And if response time would be the explanatory variable, these values could be [0.6,0.8,1.4,1.2,0.7] and [1.3,0.5,0.9,1.2,0.6,1.4].

We deal with the unknown probability distribution *f*(*X*_*r*_) by drawing from a known *conditional* probability distribution that is derived from *f*(*X*_*r*_). The general idea behind this approach is to construct a conditional probability distribution that resembles the randomization distribution that is used in a randomized experiment. We will describe two approaches, one that involves conditioning on an “informative support”, and another one that additionally involves conditioning on sufficient statistics.

#### Conditioning on an informative support

A support is the set of values that *X*_*r*_ can take (i.e., its realizations). For the purpose of producing a sensitive statistical test, we construct pairs of realizations that are maximally different from each other but have the same probability. For example, in an experiment in which accuracy is the explanatory variable, this could be such a pair: {[0,0,1,1,0],[1,1,0,0,1]}. The important feature of this pair is that, for all events with a correct response in the first realization, there is an incorrect response in the second realization. Such a pair is called an informative support, and when drawing from the permutation distribution, we condition on it. This implies that, if one of the two realizations in the informative support is actually observed, to construct the permutation distribution, we randomly select one of the two. Next, we combine it with the biological data 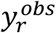 to calculate the contribution of the r-th participant to the test statistic *S*(*y*^*obs*^, *x*^*obs*^). This is almost identical to the way we construct the randomization distribution for a within-participants manipulation of the explanatory variable: instead of randomly selecting one of the two complementary condition orders, we now randomly select from the informative support.

This approach also applicable to continuous explanatory variables like RT, pupil diameter or some EEG-parameter (e.g., occipital alpha power). Assume that the observed explanatory variable is [2.7,9.3,4.9,7.5,3.1] and the interest is in a monotone relation between the biological data and this explanatory variable. For an interest in a monotone relation, an informative support can be based on order statistics: {[2.7,9.3,4.9,7.5,3.1],[9.3,2.7,4.9,3.1,7.5]}. The order statistics of the second variable ([5,1,3,2,4]) is the reverse of the order statistics for the first pair ([1,5,3,4,2]). We now describe our approach more formally.

In general, we draw from 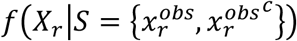, in which *S* denotes the support of the probability distribution. Here, we constrain the support to a set of two possible realizations: 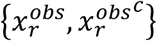, the observed condition order 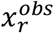 plus a condition order 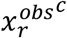 that is maximally different from the observed one. “Maximally different” is operationally defined by the researcher with the objective of maximizing the sensitivity of the statistical test. Maximizing sensitivity implies that 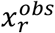 and 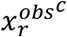 must be as different as possible with respect to the effect of interest. To prevent bias, these pairs must be chosen independently from the biological data 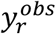. A set 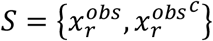 that is chosen in this way is called an informative support. A draw from the conditional probability distribution 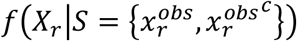 approximates a draw from a randomization distribution of which the possible realizations are as different as possible with respect to the effect of interest.

With the resulting permutation test, we test the null hypothesis of statistical independence between *X*_*r*_ and *Y*_*r*_ given that *X*_*r*_ belongs to an informative support *S*:

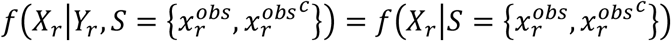

The conditioning on the informative support is crucial, because it determines the sensitivity of the statistical test: without this conditioning, the generality of the inference is low, because the outcome of the test (reject or accept) may be dominated by one or a few participants. By conditioning on an informative support, a small p-value can only be obtained if the evidence provided by the different participants is consistent along the dimension that is implied by the difference between 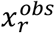 and 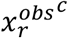. This increases the generality of the inference.

The conditional probability distribution 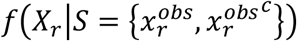 is a Bernoulli distribution parameterized by the conditional probability of observing 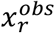 given that one either observes 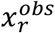 or its complement 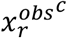:

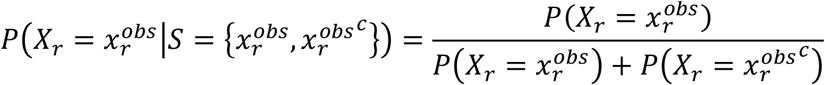

In order to draw from 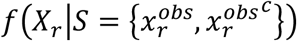, the probabilities 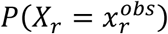 and 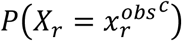 must be known. However, because *X*_*r*_ is not under experimental control, its probabilities are unknown. To deal with this problem, besides conditioning on an informative support (which determines the sensitivity of the statistical test), we will also condition on sufficient statistics.

#### Conditioning on an informative support and sufficient statistics

In this approach, we draw from the conditional probability distribution 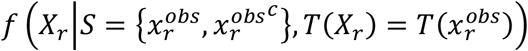, in which *T*(*X*_*r*_) is a sufficient statistic of *X*_*r*_. A sufficient statistic is a function of a random variable (here, *X*_*r*_) with the property that, by conditioning on its value, the resulting conditional probability distribution of the random variable becomes independent of the unknown parameters. Therefore, this conditional distribution only depends on the observations 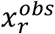 or 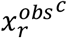, which are known. For a given probability distribution *f*(*X*_*r*_), there are multiple sufficient statistics, and these are usually specific for a particular parametric model for *X*_*r*_ (e.g., mean and variance for a normal distribution). Because it is generally unknown which parametric model governs a particular *X*_*r*_, we opt for a nonparametric sufficient statistic, the unordered data {*X*_*r*_}. For example, if 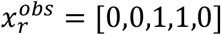, then 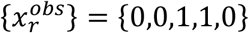. Now, given that we also condition on the informative support 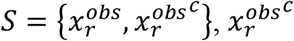 must have the same sufficient statistic as 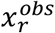, since otherwise 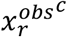 has a zero probability under 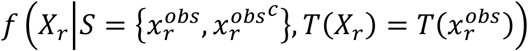. Therefore, we set 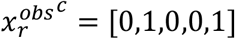 instead of 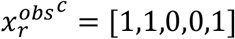. With this definition of the informative support 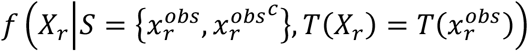 is a Bernoulli distribution with probability 0.5 for both 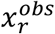 and 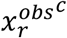.

The unordered data {*X*_*r*_} are not always a sufficient statistic. Specifically, they are not a sufficient statistic if the elements of *X*_*r*_ (the event-specific explanatory variables) have different probability distributions (e.g., their mean increases with the event number) or if they are statistically dependent (e.g., auto-correlated with some nonzero lags). To deal with such patterns in the data, one can build a parametric probability model for *X*_*r*_ that captures these patterns (e.g., a logistic regression model with a linear trend and pair-wise statistical dependencies) and condition on the sufficient statistics for this model. Parametric probability modelling also provides diagnostic tools to identify such trends and statistical dependencies.

#### The permutation test controls the FA rate

Under the permutation distribution, a p-value is calculated, which subsequently is used to take a decision about the null hypothesis of statistical independence between *X*_*r*_ and *Y*_*r*_. This permutation test controls the FA rate, and we can prove this in the same way as shown previously for observational studies with a between-units manipulation of the explanatory variable. Specifically, this proof involves the following two steps:

1. The permutation test controls the FA rate conditionally given 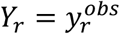, 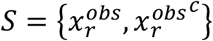 and 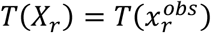, for *r* = 1, …, *n*.
2. FA rate control conditionally given 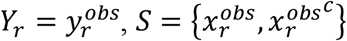 and 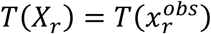 implies unconditional FA rate control.

It is important to note that, in many observational studies, not only the condition orders are not under experimental control, but also the event times *E* and the way they are partitioned in the condition-specific sets *E*_1_ and *E*_2_. In these studies, *E*_1_ and *E*_2_ are random variables, and this status has to be taken into account when proving the FA rate. Specifically, in this proof, we must condition on the realizations of *E*_1_ and *E*_2_. As shown in the other proofs for the permutation test, conditioning on random variables does not affect the FA rate control.

### Using a Permutation Test in a Randomized Study to Allow for a More General Inference

When we discussed randomization tests for studies with a within-participants manipulation of the explanatory variable, we pointed out that the generality of inference depends on the randomization distribution, and in particular, on its number of possible realizations. We now consider a randomized study in which we have used a randomization distribution *f*(*X*_*r*_) that, in retrospect, does not allow for the desired generality of inference. For instance, the randomization distribution may generate condition orders by randomly drawing from a group of condition labels with equal numbers for the two experimental conditions A and B. However, the objective of the study may be only to evaluate the effect of the two experimental conditions A and B, and this could be better evaluated by comparing one condition order with its complement. Fortunately, the permutation distribution 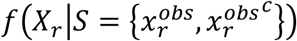 (for *r* = 1, …, *n*) allows for this. Specifically, because of the symmetry in the way the random condition orders are generated, every observed condition order 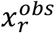 has a complement 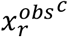 with the same probability. Now, by using the conditional distribution 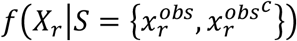 instead of the unconditional randomization distribution *f*(*X*_*r*_), we now evaluate a null hypothesis that allows for a more general inference:

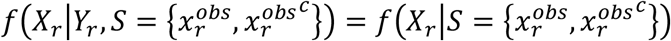

## Discussion

### Summary of the Main Contributions

We have described a general approach to nonparametric statistical testing that starts from the null hypothesis of statistical independence between the explanatory and the biological variable. We have provided a formal proof of the FA rate control under this approach. Crucially, FA rate control is achieved without auxiliary assumptions and does not depend on asymptotic arguments (e.g., increasing sample size or cluster-defining threshold). In this respect, it outperforms the existing parametric tests and some existing nonparametric tests (i.e., the sign-flipping test). Our approach is a very general one because it can be used for (1) both single- and multi-participant studies, (2) studies with a between- as well as a within-participants manipulation of the explanatory variable, (3) studies with a randomized as well as a natural explanatory variable (possibly involving between-participant differences in event times), (4) studies with a categorical as well as a quantitative explanatory variable, and (5) for testing null hypotheses that are of a fixed or a random effect type.

### Random versus Non-random Assignment to the Conditions, and Another Null Hypothesis

For the randomization tests presented in this paper, it is required to have a random explanatory variable with a known probability distribution. This has the simple but important consequence that the statistical testing procedure already starts before the data collection, namely when the units (participants) are randomly assigned to the conditions (condition orders). The result of this random assignment is then stored, and reused when the randomization p-value is calculated.

The requirement of random assignment follows from the null hypothesis of statistical independence between the biological data and the explanatory variable: this hypothesis can only be formulated for random explanatory variables. However, there is at least one other useful nonparametric null hypothesis that does not require the explanatory variable to be random: equality of the probability distributions between conditions or condition orders (see further). Both this other null hypothesis and its statistical test (see further) are closely related to the null hypothesis of statistical independence and the nonparametric tests that are described in this paper. This other null hypothesis was formulated in our earlier work (Maris & Oostenveld, 2007), and can be tested in studies in which units are non-randomly assigned to a condition or a condition order (e.g., alternating between the different condition orders).

The alternative null hypothesis of interest is the following: equality of the probability distributions of the biological data of a randomly sampled unit (*Y*_*r*_) in the different conditions (condition orders). Formally, for a study with a between-units manipulation of the explanatory variable, this null hypothesis can be written as follows:

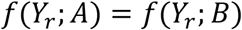

In this equation, *f*(*Y*_*r*_; *A*) and *f*(*Y*_*r*_; *B*) denote the probability distributions of the biological data in the conditions *A* and *B*. And for a study with a within-unit manipulation of the explanatory variable and condition orders ABBABABA and BAABABAB, this null hypothesis can be written as follows:

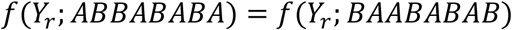

This null hypothesis of equal probability distributions can be tested by means of a permutation test, but instead of permuting the realizations of the explanatory variable (while keeping the biological data fixed), we now permute the unit-specific component data structures (while keeping the non-random explanatory variable fixed). Under the null hypothesis of identical probability distributions in different conditions, every permutation of the component data structures has the same probability. Exactly the same p-value results from these two procedures (permuting the explanatory variable and permuting the biological data), and the only difference is the null hypothesis.

In the present paper, we deliberately focused on randomization tests of the null hypothesis of statistical independence. This is because the nonparametric tests of the alternative null hypothesis (equal probability distributions) are less general. Specifically, these nonparametric permutation tests cannot be used in most event-related fMRI studies, such as a single-participant study or a multi-participant study with a within-participants manipulation of the explanatory variable.

### The Generality of the Framework

The framework presented in this paper is general, but can still be extended. Specifically, it can be extended to (1) other types of explanatory variables, (2) null hypotheses about the incremental effect of one explanatory variable given another (confounding) explanatory variable, and (3) null hypotheses about interaction effects. Starting with the first point, one must first observe that the null hypothesis of statistical independence is not restricted to a particular type of explanatory variable (two or more levels, nominal or quantitative, uni- or multivariate). The test statistic, however, does depend on the type of the explanatory variable. Specifically, for nominal explanatory variables with more than two levels, an F-statistic is typically used, and for quantitative explanatory variables a regression coefficient t-statistic. And when the explanatory variable is multivariate, one typically uses a test statistic based on a canonical correlation (e.g., Wilks’s lambda).

Second, our framework can deal with incremental effects of one explanatory variable (*X*_1_) given another (confounding) explanatory variable (*X*_2_). Incremental effects are highly relevant in observational studies with correlated explanatory variables, of which one is of interest and the other(s) is (are) confounding. For example, one may be interested in the association between disease status (e.g., diseased vs. healthy) and some biological variable, but the control group cannot be matched to the disease group on all possible variables. Now, if there is an age difference between the two groups, age could be a confounding variable in the association between disease status and the biological variable.

In our framework, we formulate the hypothesis of no incremental effect as *conditional* statistical independence (for short, conditional independence) between *Y* and *X*_1_ *given X*_2_. This can be expressed formally as follows:

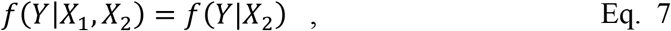

This equation formalizes the notion that, conditionally on *X*_2_, the explanatory variable *X*_1_ is not associated with the biological variable *Y*.

The permutation test for this hypothesis of conditional independence is very much along the same lines as a permutation test for the regular (unconditional) independence between *X* and *Y*. The essential difference is that the test statistic now is evaluated under a so-called *grouped permutation distribution*, in which “grouped” denotes that we have one permutation distribution for every unique value (e.g., age or age group) in the realization 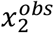 of *X*_2_, the random variable on which we condition. Specifically, we will draw the realizations of *X*_1_ (e.g., disease status) in parts, and each part corresponds to the components of *X*_1_ for which the participants all have the same realization in the vector 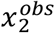 (e.g., the same age or age group). Each of these parts is drawn from its own permutation distribution, and together they form a grouped permutation distribution. Thus, the decision about the null hypothesis of conditional independence is taken on the basis of a p-value that is calculated under this grouped permutation distribution.

The FA rate of this grouped permutation test is controlled under the new conditional null hypothesis in Eq. 7. The formal proof of this FA rate control is very similar to the one for the unconditional independence between *X* and *Y*. It is again a two-step proof in which the first step proves conditional FA rate control, and the second unconditional FA rate control.

Third, our framework allows for testing null hypotheses about interaction effects. Different from the null hypothesis considered up to now, an interaction effect null hypothesis pertains to processed instead of raw data. This is most easily explained for a study with one explanatory variable that is manipulated within-participants, and another one that is manipulated between-participants. For every individual participant, we quantify the effect of the within-participants explanatory variable as the difference between condition-specific regression coefficients. We denote these participant-specific effect quantifications by *Y*_*r*_ (at the level of single participants) and *Y* (an array at the level of the whole sample of participants). The interaction effect null hypothesis is now formulated as statistical independence between the effect quantification array *Y* and the between-participants explanatory variable *X*. This null hypothesis is tested by means of a randomization or permutation test, depending on whether *X* is a randomized or a natural explanatory variable.

### Parametric versus Nonparametric Statistical Tests

In this paper, we advocate for the use of nonparametric statistical tests because of their exact FA rate control. This is a clear advantage over parametric statistical tests, whose exact FA rate control depends on auxiliary assumptions and/or on asymptotic arguments (e.g., an increasing sample size or cluster-defining threshold).

Parametric statistical tests have two valuable properties which nonparametric tests do not have, or which cannot be proved: (1) some parametric statistical tests are uniformly most powerful (UMP, Lehmann, 1986), and (2) parametric probabilistic modeling of the biological data allows for statistical testing at the level of the free parameters of the model, which is highly informative given that the model is valid. However, both properties have only a limited relevance in current cognitive neuroscience. Starting with the first, a statistical test is defined to be UMP if there is no other statistical test that is uniformly (i.e., over all values of the parameter of interest) more powerful (sensitive). Now, if the parameter of interest is a scalar expected value then, under the usual auxiliary assumptions (independence, normality, equal variance), the univariate t-statistic is UMP. However, this property does not hold for multivariate expected values and the corresponding multivariate t-statistic (Hotelling’s *T*^2^). Therefore, for scientific disciplines that collect multivariate data, such as cognitive neuroscience, the UMP-status of the univariate t-statistic is not very relevant.

Second, it would be extremely difficult to argue for nonparametric statistical testing if a valid probabilistic model of the data would be available, especially if this model would have a low number of free parameters as compared to the dimensionality of the data. Such a model would allow for statistical tests with respect to the free parameters of the model (e.g., a likelihood ratio test). In a sparsely parameterized model, this would be very parsimonious, as it would allow for an interpretation of the effect in a low-dimensional subspace of the full data space. Unfortunately, such a sparsely parameterized probabilistic model is not available for any of the common biological data collected in cognitive neuroscience.

## Conclusion

Starting from the null hypothesis of statistical independence between the explanatory and the biological variable, we have developed a general approach to nonparametric statistical testing. The formal core of this approach is a proof of its FA rate control, which is valid for a very wide range of studies (single- and multi-participant studies, with a between- as well as a within-participants manipulation, with a randomized as well as a natural explanatory variable, with a categorical as well as a quantitative explanatory variable, and for testing null hypotheses that are of a fixed or a random effect type). It is easy to extend the framework to multivariate explanatory variables, null hypotheses about the incremental effect of one explanatory variable given another, and null hypotheses about interaction effects. Although we do not claim that parametric statistical tests are now obsolete, for a number of statistical problems in cognitive and medical neuroscience, a solution can be found in this nonparametric framework.

## Notes

#### Summary of Updates

The figure references were messed up by the conversion to .pdf. I have correct this.

